# Extended Poisson Gaussian-Process Latent Variable Model for Unsupervised Neural Decoding

**DOI:** 10.1101/2024.03.04.583340

**Authors:** Della Daiyi Luo, Bapun Giri, Kamran Diba, Caleb Kemere

## Abstract

Dimension reduction on neural activity paves a way for unsupervised neural decoding by dissociating the measurement of internal neural state repetition from the measurement of external variable tuning. With assumptions only on the smoothness of latent dynamics and of internal tuning curves, the Poisson Gaussian-process latent variable model (P-GPLVM) (Wu et al., 2017) is a powerful tool to discover the low-dimensional latent structure for high-dimensional spike trains. However, when given novel neural data, the original model lacks a method to infer their latent trajectories in the learned latent space, limiting its ability for estimating the internal state repetition. Here, we extend the P-GPLVM to enable the latent variable inference of new data constrained by previously learned smoothness and mapping information. We also describe a principled approach for the constrained latent variable inference for temporally-compressed patterns of activity, such as those found in population burst events (PBEs) during hippocampal sharp-wave ripples, as well as metrics for assessing whether the inferred new latent variables are congruent with a previously learned manifold in the latent space. Applying these approaches to hippocampal ensemble recordings during active maze exploration, we replicate the result that P-GPLVM learns a latent space encoding the animal’s position. We further demonstrate that this latent space can differentiate one maze context from another. By inferring the latent variables of new neural data during running, certain internal neural states are observed to repeat, which is in accordance with the similarity of experiences encoded by its nearby neural trajectories in the training data manifold. Finally, repetition of internal neural states can be estimated for neural activity during PBEs as well, allowing the identification for replay events of versatile behaviors and more general experiences. Thus, our extension of the P-GPLVM framework for unsupervised analysis of neural activity can be used to answer critical questions related to scientific discovery.

## 1 Introduction

Memory critically requires firing of neurons in the hippocampus both during ongoing experiences and afterwards, as the resultant memories are consolidated. While rodent studies have focused on spatial memories, hippocampal neurons can be generally understood to represent the conjunction of the sensory features associated with a particular context (Moser et al., 2015), and the temporal sequences that connect local contexts across time during an experience (Eichenbaum and Cohen, 2014; Eichenbaum, 2017). Importantly, sequential firing patterns of neural ensembles reactivate in a time-compressed manner during some of the population burst events (PBEs) that occur during sharp-wave ripple oscillations in sleep or quiet wakefulness (Wilson and McNaughton, 1994; Skaggs and McNaughton, 1996; Kudrimoti et al., 1999; Nádasdy et al., 1999). By decoding those events, it has been shown the replay trajectories show a continuum of conformity to the original experience, including variability in momentum and both forward and reverse re-expression (Lee and Wilson, 2002; Foster and Wilson, 2006; Diba and Buzsáki, 2007; Csicsvari et al., 2007; Davidson et al., 2009; Krause and Drugowitsch, 2022). Traditionally, individual replay events have been identified based on a strong assumption of ordered consistency with patterns expressed during exploration. Consequently, un-ordered replay of contexts, or ordered replay of more complicated routes are often excluded from subsequent analysis. Thus, while much has been learned about memory consolidation and recall from the study of replay, existing approaches have colored our understanding. Therefore, a technique for decoding neural activity without strongly stereotyping the patterns represented or requiring a specifically spatial encoding model would be a powerful tool for understanding memory.

To extract the information from spike trains with minimal prior assumptions, one practicable approach is to find a low-dimensional embedding that can reveal the underlying dynamics. The Poisson Gaussian-process latent variable model (P-GPLVM) proposed by Wu et al. (2017) is a probabilistic, nonlinear, and dynamic dimension reduction approach. It infers temporally smooth low-dimensional latent neural trajectories and smooth, non-parametric internal tuning curves from spike trains without referring to external variables. This model consists of Poisson spiking observations and two Gaussian processes, one governing the temporal evolution of latent variables and another governing the nonlinear mapping from high-dimensional neural data to the low-dimensional latent variables.

In the learned low-dimensional latent space, (1) by mapping any possible external variable unto the embedding, how external variables are represented in this latent space can be revealed; (2) by measuring the repetition of internal neural state (relative locations to the low-dimensional training data embedding), the repetition of neural activity encoding external experiences can be detected (Yu et al., 2009; Rubin et al., 2019; Nieh et al., 2021). Unsupervised neural decoding can be achieved by dissociating the measurement of internal neural state repetition from the measurement of external variable tuning. However, an approach for inferring latent variables of new data points in the learned latent space is lacking from the original P-GPLVM model, limiting its utility for decoding.

In this paper, we extend the P-GPLVM framework to enable the latent variable inference of new inputs constrained by the learned smoothness parameters and tuning curves (Figure 1), and develop subsequent analyses to evaluate the new input data in the latent space. We also present a preprocessing pipeline for PBE decoding when using models trained from behavioral data. The original P-GPLVM can be used to reveal encoded information in neural activity and discover neural trajectory evolution but only within the given training data. This extended model and PBE preprocessing pipeline, especially, makes use of information learned in the model and enables the effective and unsupervised decoding of new neural activity both during behavior and during PBEs.

**Figure 1:**
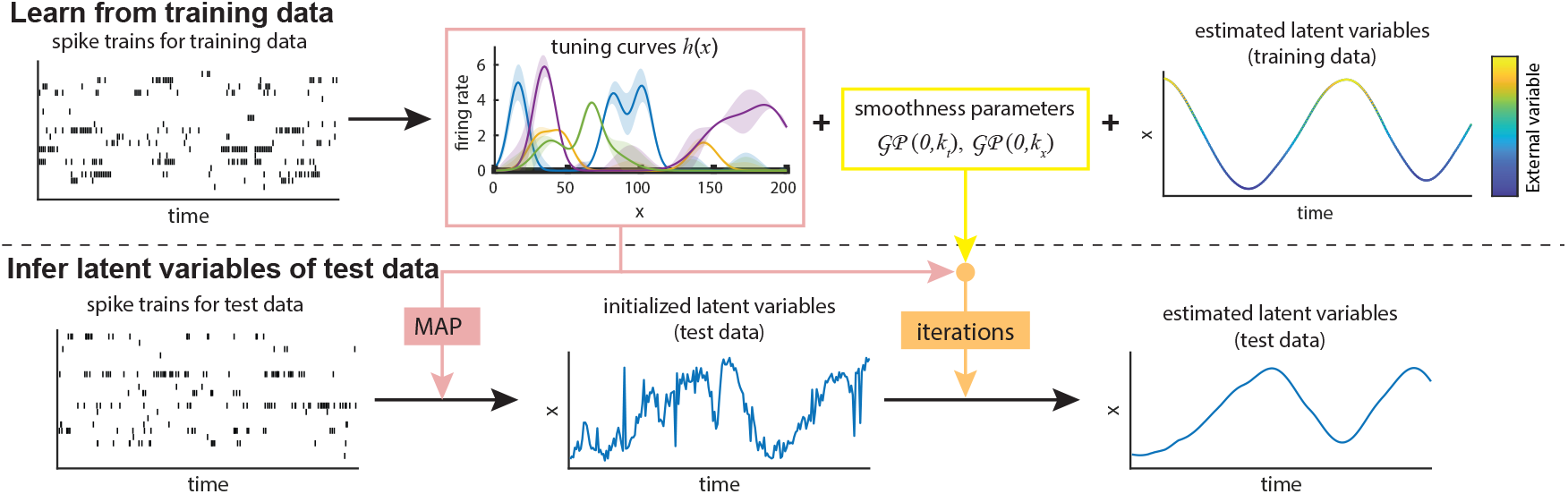
Schematic diagram of the extended P-GPLVM model. Internal tuning curves and smoothness information learned from training data are then used to constrain the inference of test data latent variables in the same latent space.

## 2 Poisson Gaussian-process latent variable model

### 2.1 Model structure

Binning spike counts into *M* time bins from *N* neurons creates the matrix of spike trains 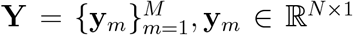. The *m*^*th*^ time bin was recorded at time *t*_*m*_, *m ∈* (1, *· · ·, M*). In this model, two latent variable matrices will be learned: the log firing rate for Poisson spiking, 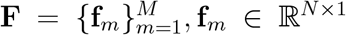, and the *P* -dimensional latent variables, 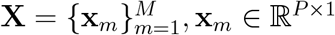.

#### Latent dynamics

Each feature of the latent variable matrix **X** evolves according to a Gaussian process depending on time *t, x*_*p*_(*t*) *∼ 𝒢 𝒫* (0, *k*_*t*_), *p ∈* (1, *· · ·, P*), where *k*_*t*_(*t, t*^*′*^) = *r* exp(*−*|*t − t*^*′*^|*/l*), governing the temporal smoothness of *x*_*p*_. Writing as a multivariate normal distribution,

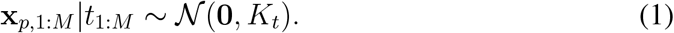

The covariance *K*_*t*_ is an *M × M* matrix with entries *k*_*t*_ at all pairs of time bins.

#### Nonlinear mapping

Let *h* : ℝ^*P*^ *→* ℝ be a nonlinear mapping function, describing the firing rate of a neuron in the *m*^*th*^ time bin as *λ*_*m*_ = *h*(**x**_*m*_). The log tuning curve of the *n*^*th*^ cell in response to the latent variable **x** is modeled as another Gaussian process as *f*_*n*_(**x**) = log *h*_*n*_(**x**) *∼ 𝒢 𝒫* (0, *k*_*x*_), *n ∈* (1, *· · ·, N*). This process has 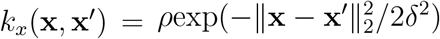 as a spatial covariance function. Therefore, the log firing rates of the *n*^*th*^ neuron in all time bins have the multivariate normal distribution

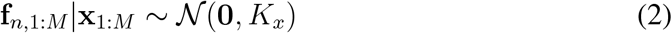

with the *M × M* covariance matrix *K*_*x*_, whose entries are *k*_*x*_ evaluated for all pairs of **x**. By combining **f** over all the neurons, **F** *∈* ℝ^*N×M*^ as firing rates in units of spike counts per time bin are obtained. *Note that this is per* bin, *rather than per* second. *This has consequences for PBE decoding, as described below*.

#### Poisson spiking

Finally, for the *n*^*th*^ neuron in the *m*^*th*^ time bin, observed spike counts *y*_*n,m*_ are assumed to be drawn from a Poisson process given the latent firing rate *λ*_*n,m*_ = exp(*f*_*n*_(**x**_*m*_)),

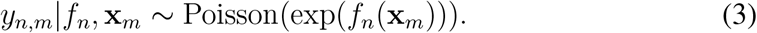

### 2.2 Model training

During training, the model iteratively infers latent variables of training data without mapping constraints and optimizes smoothness parameters. In each iteration, **X** is first fixed and **F** is optimized. For each neuron, the posterior over **f**_*n*_ is given by

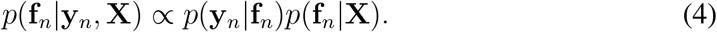

The optimal 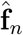 is found by maximizing the log conditional distribution

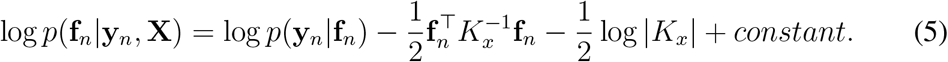

Then, fixing **F**, the optimal **X** is discovered by maximizing the conditional likelihood

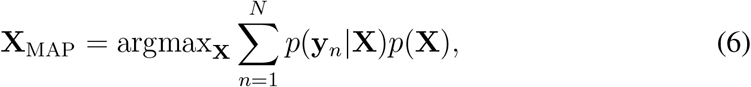

where *p*(**y**_*n*_|**X**) is given by

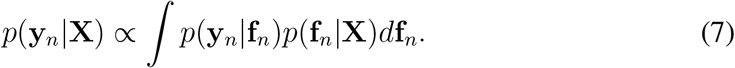

Using Laplace’s method, the approximated log likelihood conditioned on **X** is

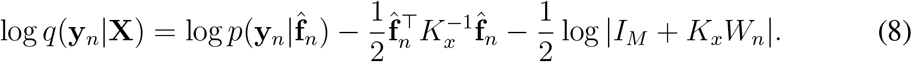

where *W*_*n*_ = *−∇∇* log *p*(**y**_*n*_|**f**_*n*_). Hyperparameters ***θ*** = *{ρ, δ, r, l}* are found by maximizing the same likelihood function.

## 3. Extensions of P-GPLVM

### 3.1 Constrained latent variable inference of new data

#### Latent variable initialization

During model training, P-GPLVM learns smoothness parameters, θ, and internal mapping function, log *h*_*n*_ : ℝ^*P*^ → ℝ, parameterized by latent variables 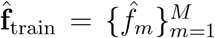 for each cell and 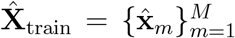 from training data. In the original P-GPLVM paper, the tuning curve vectors **f**_grid_ are evaluated at the grid of latent variables 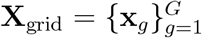 using a joint Gaussian distribution with 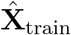 and 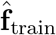.

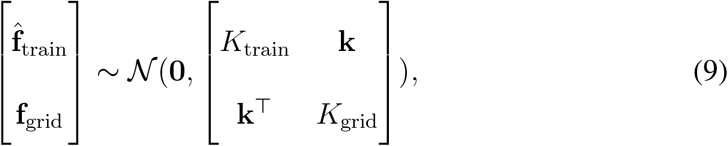

where 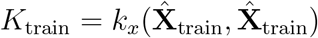, *K*^grid^ = *k*^*x*^ (**X**^grid^, **X**^grid^), and 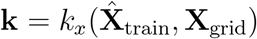, are the covariance matrices. The entry 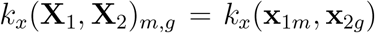. The posterior distribution of **f**_grid_ can then be written as

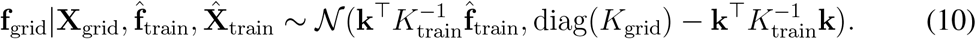

Given new input data from the same group of neurons, 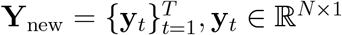, the latent variables 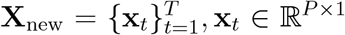, are initialized with elements in **X**_grid_ that maximize the posterior probability using a Bayesian approach. At the *t*^th^ time bin, the posterior over each element in **X**_grid_ is

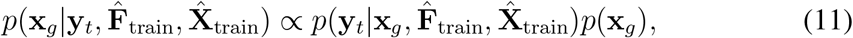

where we’ve assumed a uniform prior over **x**_*g*_ during initialization. The prior over **y**_*t*_ is

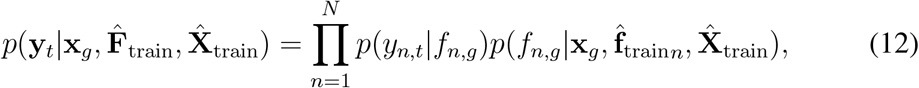

The latent variable **x**_*t*_ corresponding to **y**_*t*_ is initialized with the **x**_*g*_ with maximum posterior.

#### Impose smoothness constraints

Merely using MAP tuning curves to constrain the latent variable inference of new input data omits the learned temporal smoothness in the latent space. To impose both temporal smoothness and tuning constraints, subsequent iterations are needed. We noted that employing **f**_grid_ and **X**_grid_ as tuning curve vectors is computationally expensive – considering *a* samples per dimension, a *P* -dimensional **X**_grid_ contains *a*^*P*^ elements. Moreover, a majority of the grid elements contain minimal information as no 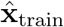 locate nearby. To save computational costs during iterations and to preserve the tuning details of 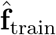 in response to 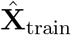 as much as possible, 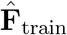 and 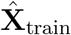 were directly used as the tuning curve vectors (TC), 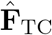 and their encoded latent variables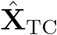.

Iterations are similar to those during unconstrained inference as described in Section 2.2, but the smoothness parameters are fixed as learned and the mapping function log *h*_*n*_ is constrained by 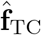 and 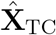. This is achieved by substituting 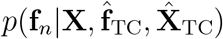 for *p*(**f**_*n*_|**X**) in Eq. 4 and 7. As in Eq. 9, the joint distribution with 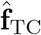 and 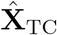 gives posterior of **f**_*n*_ as

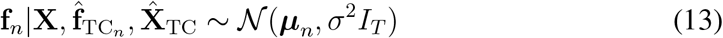

where 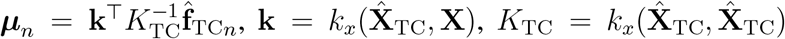 and *σ*^2^ is the observation noise. Correspondingly, Eq. 5 is modified as

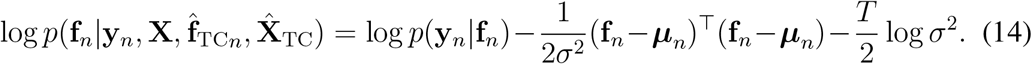

Finally, obtaining the optimal 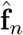, Eq. 8 is modified as

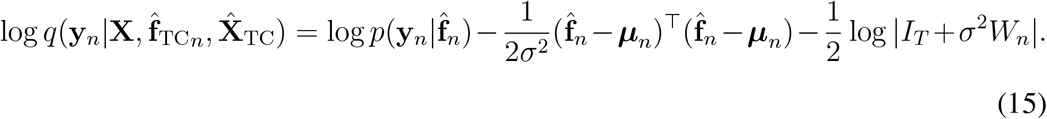

### 3.2 Preprocessing for PBE decoding

Critically for our model, neural activity during PBEs is temporally-compressed relative to that expressed during exploration. Consequently, the firing rate model learned on exploration-related neural activity will not properly model PBEs. This is addressed in two steps, choosing a shorter time bin (implicitly compressing time) and explicitly scaling the model parameters. Temporal compression implies that the time intervals between pairs of place cells activating during exploration running are expected to be proportional to those during replay. Thus, the cross-correlation histogram (Harrison et al., 2013; Karlsson and Frank, 2009) of spike trains from place cell pairs is used to find the PBE time bin size that would best match the binned behavioral data. Place cells are identified in either running direction and then are pooled. For each place cell pair with spike trains *s*_1_, *· · ·, s*_*M*_ from one cell and *t*_1_, *· · ·, t*_*N*_ from another, time lags between any pairs of spikes (*s*_*m*_, *t*_*n*_) within 4 seconds are recorded. The histogram of all these spike time lags is the cross-correlation histogram (CCH) for this cell pair. All histograms are centered at 0s time lag and share the same number of bins. For ease, the histogram bin size of CCH during active exploration is set as the same as that used for P-GPLVM analysis of this data. Among all pairwise combinations of place cells, cell pairs with place field (where peak firing rate occurs) distances larger than the 80^th^ percentile of the distribution of travel distance in 4 seconds are excluded from subsequent analysis. For each included cell pair, the Pearson correlation between CCH during exploration and CCH during PBEs is evaluated. The optimal bin size for PBEs is found by maximizing the sum of all significant positive Pearson correlation coefficients using a range of possible PBE bin sizes.

As the unit of our tuning curves is spike counts per time bin, for PBE-decoding, the firing rates of P-GPLVM tuning curves learned using neural activity during active exploration must be further scaled. Scaling by the ratio of the time bin sizes is found to yield good PBE-decoding performance.

### 3.3 Analysis in latent space

#### 3.1 Number of well-separated manifolds

The hippocampus “remaps” between different environments, meaning that whether a given neuron is active, and if so, how its spatial tuning will relate to that of other neurons is essentially random between different environments (Moser et al., 2015; Alme et al., 2014) (though recent studies have challenged this concept (Cai et al., 2016)). This implies that different environments should correspond to well-separated manifolds in latent space. This idea intuitively leads to the investigation that if the low-dimensional latent variable 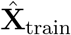 is organized in a single manifold or in multiple well-separated manifolds. This procedure consists of constructing a *K*-nearest neighbor graph in 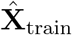 and then counting the number of connected components in the graph. Components containing less than 3% of total number of points are considered ”residual components”. Starting with a small number, *K* is gradually increased until there are no residual components left. The number of well-separated manifolds will stabilize as *K* continues to increase for a multiple times. This stabilized number is defined as the number of connected components in 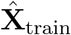. Each separated manifold is denoted as 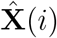 hereafter.

#### 3.2 Congruence with the learned manifold(s)

In the latent space, similar trajectories indicate repetition of neural state encoding similar external experiences. To assess new test data, whether or not their latent variables have similar spatial distribution and dynamics to those of the training data is investigated. Training and test data can contain multiple continuous recording segments. Each continuous segment is a single neural trajectory in the latent space. When the neural patterns and dynamics of new inputs **Y**_new_ match the learned model perfectly, their estimated latent trajectories **X**_new_ will (i) span the manifold of 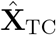 and (ii) progress smoothly along it. Three measures are proposed to evaluate congruence of new input data with a learned model.

##### Log likelihood

We use log likelihood to measure how well the learned P-GPLVM model fits the test data. The joint probability of **Y, F, X** for test data is computed as

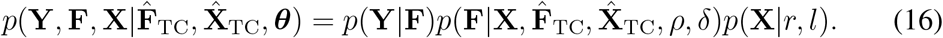

The log likelihood (LLH) of a continuous segment in test data is written as

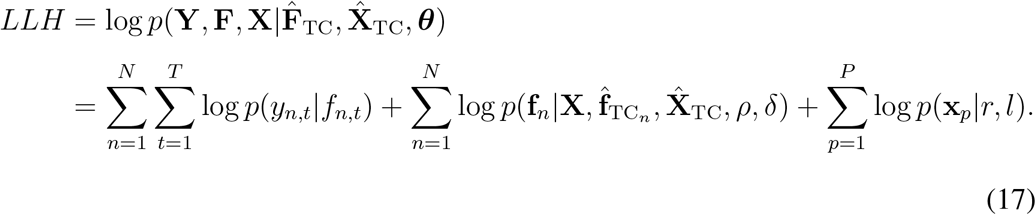

Recall that

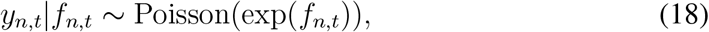

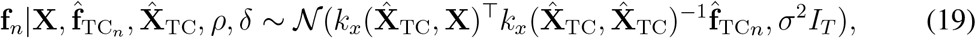

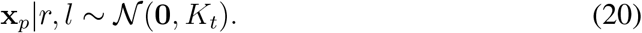

##### Spatial consistency

This metric measures how well a latent trajectory, 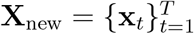, spans the latent variable manifold(s) of the tuning curve vector,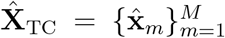. Firstly, for each point **x**_*t*_ in the trajectory, its *K*-nearest neighbors are searched in the manifold 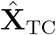, obtaining the set of its nearest neighbors 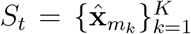 and its distances to those neighbors 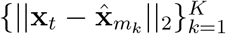. When the trajectory spans the manifold nicely, the union of neighbor sets, 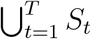, should have much more elements than by chance. The number of all identified neighbors is then weighted by the neighbor distances to integrate the neighborhood quality into the measure.

For each 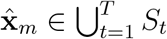, its distance to its closest point in 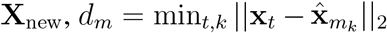 is estimated. Then, the spatial consistency of this trajectory in the latent space is computed as

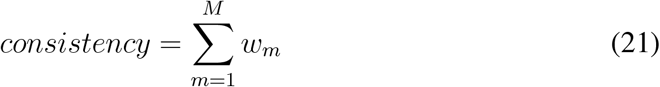

where

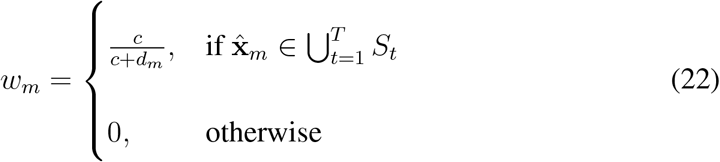

which means, when one 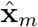 is identified as a neighbor, it contributes to the spatial consistency. This contribution is weighted by the distance to its nearest neighbor on the trajectory, which is 1 when the distance is 0, and then decay as the distance increases, reaching at 0.5 when the distance is *c*. The parameter *c* is set as 20% of the standard deviation of 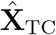. When 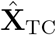 contains more than one separated manifold, the standard deviation is computed as the average within-manifold standard deviation:

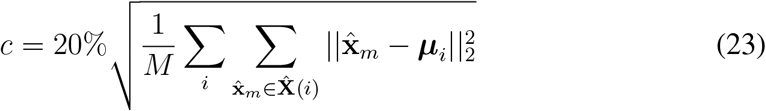

where ***μ***^*i*^ is the center of manifold 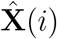.

Higher value of spatial consistency indicates that this trajectory travels a long way along the tuning curve manifold(s) rather than only jiggling at a small local area or locating far away. The spatial consistency to 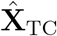 is first used to evaluate the neural trajectory behavior in the latent space. Further, when 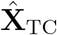 contains more than one well-separated manifold 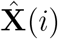, we also estimate the manifold contribution ratio to the consistency value, which is computed as in Eq. 21 but only for 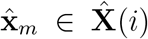. The ratio between these contribution values from manifolds indicate to which manifold the trajectory belongs.

##### Average step distance

This metric reveals the consistency between temporal adjacency and pattern-matched location adjacency. The average Euclidean distance of each step in each latent trajectory of **X**_new_ is estimated. In general, the tuning curve constraint tends to drive **X**_new_ towards their pattern-matched locations on the 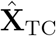 manifold. When the new data have different temporal sequences from the training data, pattern-matched locations of temporally adjacent points are mostly distant, leading to longer step distances.

## 4 Experiments and results

The dataset used in this paper is a neural recording collected in the dorsal hippocampal CA1 and CA3 areas from a rat. The recording started with a 3 hours rest session (pre), and then the animal was exposed to a novel maze (maze1) for 50 minutes, running back and forth to get rewards at the two ends. Following another two hours rest (post1), the animal explored another novel maze (maze2) for 50 minutes. Lastly, the animal went through a 5 hours rest session (post2). Spike trains during the two maze exploration sessions were binned into 500 ms time bins. Animal positions were linearized and then running periods (speed*>*3cm/s and peak speed*>*5cm/s) were extracted along with their corresponding time bin indices.

### 4.1 Exploration in one maze

We first examined this P-GPLVM with the neural activity when the animal was exposed to only one maze (maze1). The linearized animal positions during running in maze1 exploration session are shown in Figure 2A and corresponding original positions in Figure 2B. One chunk of spike trains was selected as training data to train the model (orange lines in Figure 2A, colored scatters in Figure 2B) and then the spike trains of a continuous running segment in maze1 was used as test data to evaluate the decoding performance (black line in Figure 2A,B).

**Figure 2:**
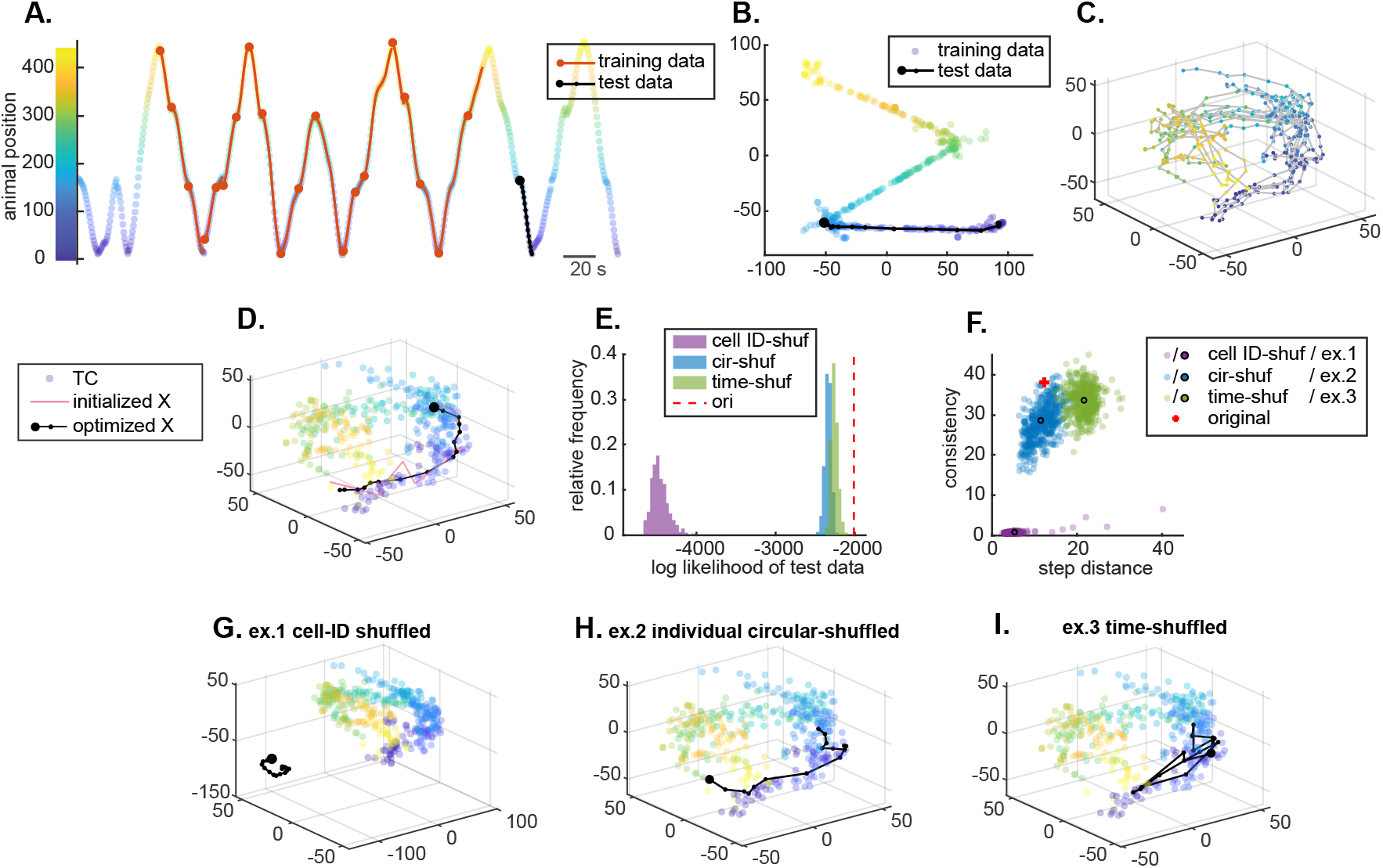
**(A)** Linearized animal position during running periods in maze1 exploration session. The orange / black lines indicate the segments used as training / test data. The starting of each continuous segment is marked by a big filled circle. Each step in the test data is marked by small dots. Scatters are color-coded by linearized animal positions, same in subsequent panels (B-D, G-I). **(B)** Colored scatters of original animal positions in training data. The black line indicates the same test data segment in (A). **(C)** Latent variables of training data in the learned P-GPLVM. Continuous segments are connected by lines as latent neural trajectories. **(D)** Inferred neural trajectory of test data in the learned latent space. The semi-transparent markers are identical with the dot scatters in (C), which, in this panel, indicates the latent variable manifold of tuning curve vectors, 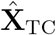. The black line depicts the inferred neural trajectory of test data, progressing along the manifold 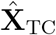 in accordance with the real animal positions (from blue to purple). **(E)** Log likelihood of the original test data (dashed red line) and its shuffled versions (histograms). **(F)** Estimated step distances vs. spatial consistency to 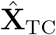 of the inferred neural trajectory in (D) compared with its cell ID-shuffled, individual circular-shuffled, and time-shuffled versions. The three black-edged circles indicate one example of cell ID-shuffled test data (ex.1), of individual circular-shuffled test data (ex.2), and of time-shuffled test data (ex.3), respectively. **(G, H, I)** Same presentation as in (D), but the inferred latent trajectories of ex.1-3 in (F).

After fitting the model, smoothness parameters ***θ*** and latent variables of training data, 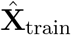 and 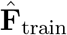, were obtained. Only one well-separated manifold is found in 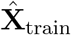, suggesting that there is no multiple distinctly different neural processes found in this training data. Given the known spatial tuning in hippocampus, to examine the information encoded in the latent space, 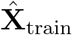 are color-coded with the corresponding animal linearized positions. Position information appears to be encoded along the manifold smoothly (Figure 2C). The latent neural trajectories seem to be a bit unkempt, which is still reasonable because this was the animal’s first exposure to maze1.

Next, we estimated the internal state repetition of the test data in the learned latent space. In Figure 2D, the manifold of tuning curve vector latent variables 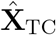, which is 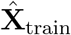 in this setting, is shown as the colored semi-transparent markers. Using the Bayesian approach, the initialized latent variables of test data appears as a bumpy trajectory (light red line). Applying the constrained inference, the latent variables converged to a smooth latent trajectory after 15 iterations (black line). By comparing the color in Figure 2B and Figure 2D, it is obvious that, the inferred neural trajectory of test data proceeds closely along those of training data with similar experiences, where the animal traveled from positions in blue to positions in purple, in accordance with the repetition of similar internal neural state.

To validate this internal neural state repetition and contrast the inferred latent variables of test data, three kinds of surrogate data were generated:

- Cell identity(ID)-shuffled data, by randomly permuting the cell identity of the test data, which scrambles the original neural coactivation patterns and permutes neuronal firing rates, but preserves the temporal sequences.
- Local individual circular-shuffled (cir-shuf) data, by randomly and circularly permuting the test data of each cell individually within each continuous segment, which scrambles coactivation patterns but preserves the temporal smoothness and local neuronal firing rates.
- Local time-shuffled data, by randomly permuting the temporal order of the test data within each continuous segment, which preserves the coactivation patterns but scrambles the time sequences.

For each shuffle type, we generated 500 surrogate test data. Metric values and example latent trajectories of the original and surrogate data are shown in Figure 2.

Figure 2E depicts the distribution of LLH values of the inferred latent variables of original test data and of the surrogate test data. Overall, the original test data has a higher LLH value than all the surrogate data. Figure 2F shows their step distance and spatial consistency values in scatters. Inferred latent trajectories of example surrogate data are shown in Figure 2G-I, whose metric values are indicated in the black-edged scatter in Figure 2F. In the latent space, the trajectories of cell ID-shuffled and of time-shuffled test data look distinct from that of the original test data trajectory, while the circular-shuffled test data trajectory has visually similar behavior.

Note that, the iterative inference process searches for an optimal balance between temporal smoothness and tuning curve constraints. Because of the changed relative firing rates among cells in cell ID-shuffled data, the tuning curve constraints push the trajectories far away from 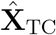, where temporal smoothness constraint dominates, leading to small step distances, diminutive spatial consistency values, and the lowest LLHs among all types of shuffled data. With preserved local neuronal firing rates, circular-shuffled data trajectories stay in the similar area as the original data, but due to the scrambled coactivation patterns and preserved temporal smoothness, the tuning curve constraints are compromised by the temporal smoothness constraint, resulting in less likely but smooth trajectories with step distances comparable to the original test data, slightly lower spatial consistency values, and significantly lower LLH values. As for time-shuffled data, with intact coactivation patterns at each time point, the tuning curve constraints are dominating during the inference, driving the latent variables to the corresponding locations on the manifold of 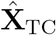 as the original test data does, while the temporal smoothness is sacrificed, resulting in spiky trajectories with large step distances, comparable spatial consistency, and slightly lower LLH values.

We contrast the original test data with its shuffled versions using a modified z-scored LLH, calculated by dividing the difference from the median by the median absolute deviation of shuffled data LLH. The z-scored LLH of the original test data among its cir-shuf versions is 13.56, and among its time-shuffled versions, 8.98, indicating that the test data is a valid sequential repetition of internal neural states captured in the training data. This measure separately verifies the matching of coactivation patterns between test data and tuning curves (when compared with cir-shuf test data), and the matching of temporal sequences (when compared with time-shuffled test data).

### 4.2 Exploration in two mazes

Next, we asked if our approach can still work well for neural activity during exploration in more than one environment context. One chunk of neural data during running in maze1 (orange lines in Figure 3A, colored scatters on the left in Figure 3B) and another in maze2 (blue lines in Figure 3A, colored scatters on the right in Figure 3B) are selected and concatenated as the training data to train a new P-GPLVM. The test data is from maze1, the same as in the previous section (black line in Figure 3A, B). All the semi-transparent markers in the scatter plot are colored by the corresponding linearized animal positions.

**Figure 3:**
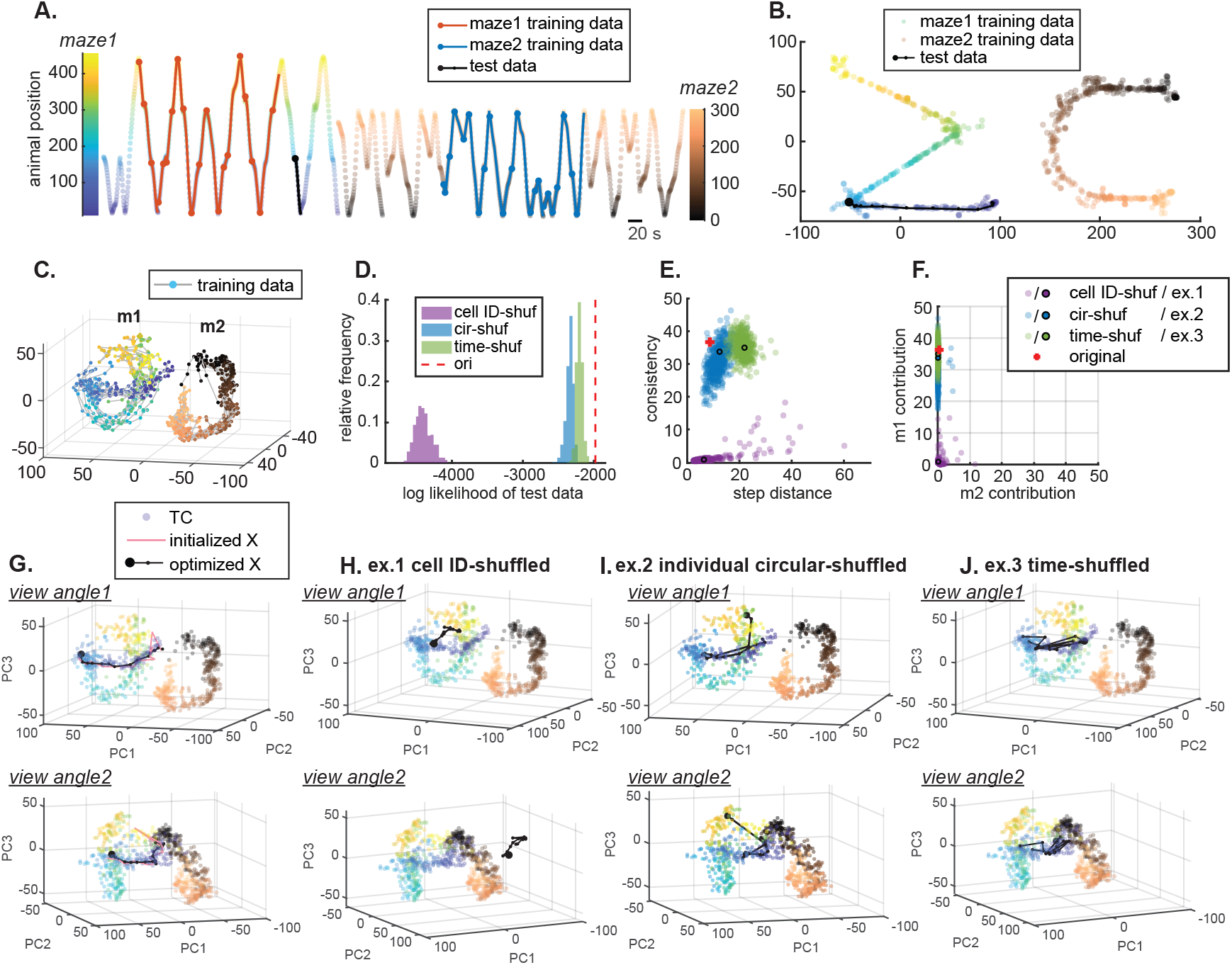
**(A)** Linearized animal position during running periods in maze1 and maze2 exploration sessions. The orange / blue lines indicate the segments in maze1 / maze2 used as training data. The black line indicates the test data. Scatters are color-coded by contexts and linearized animal positions, same in (B-C, G-J). **(B)** Scatters of original animal positions in training data from maze1 (left) and maze2 (right). The black line indicates the same test data segment as in (A). **(C)** Following fitting the training data, latent variables are separated into two distinct manifolds, m1 and m2. Each data point is colored by its corresponding linearized animal position as in (A). **(D)** Log likelihoods (LLH) of test data (red dashed line), cell ID-shuffled (purple histogram), individual circular-shuffled (blue histogram), and time-shuffled surrogate data (green histogram). **(E)** Inferred latent neural trajectory step distance vs. spatial consistency to 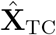 of original test data (red +), of cell ID-shuffled (purple scatters), of individual circular-shuffled (blue scatters), and of time-shuffled (green scatters) surrogate data. Example surrogate data indicated in one black-edged circle from either shuffle type is then shown for visualization. **(F)** Manifold contribution to neural trajectory spatial consistency from m1 vs. from m2, of original and surrogate test data. **(G)** Latent neural trajectories as initialized (light red line) and optimized after iterations (black line) of test data in the latent space. Latent variables of tuning curve vectors, 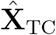, are shown in semi-transparent markers color-coded by the animal positions, same as in (C). Both initialized and optimized trajectories of test data are associated with the corresponding context manifold. Unlike the bumpy initialization, optimized neural trajectory is smooth and progresses according to the animal positions. **(H-J)** Same representations as in (G), inferred trajectory of **(H)** ex.1 for cell ID-shuffled test data, **(I)** ex.2 for individual circular-shuffled test data, and **(J)** ex.3 for time-shuffled test data, as indicated in black-edged circles in (E,F).

In 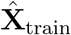 of the newly learned model, two well-separated manifolds, m1 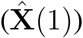 and m2 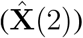, are found, each of whom turns out to contain all data points from one of the maze contexts. Using the same color code as in Figure 3A and B on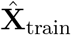, position information appears to be encoded along the corresponding context manifold smoothly (Figure 3C). Thus, this approach is capable of detecting the presence of two environment contexts as well as capturing the encoded position information merely from the neural activity in the training data.

Again, cell ID-shuffled, individual circular-shuffled, and time-shuffled surrogate test data were generated. Along with the original test data, their latent variables in the new latent space are inferred, which behave quite similarly to those in Section 4.1. Figure 3D depicts the distribution of LLH values of the original test data and the surrogate test data. Cell ID-shuffled test data have much lower LLHs than all the other type of data. Circular-shuffled data have slightly smaller LLHs than time-shuffled data. The original test data has a higher LLH value than all surrogate data, where z-scored LLH among its circular-shuffled versions is 11.75, and among its time-shuffled versions, 8.40, confirming that the test data is a valid internal neural state repetition.

Next, we investigated the content of this state repetition by looking into the latent variable locations relative to 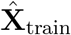. The inferred latent trajectory of the test data and its initialization in this new latent space are shown in Figure 3G. The trajectory is similar to those neural trajectories of training data with similar experiences, traversing the same arm of maze1, from positions blue to purple just as in Figure 3B.

As a high-level summary of the spatial distribution and dynamics of these neural trajectories, Figure 3E shows the step distance and spatial consistency values to 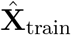, while Figure 3F shows the manifold contribution to spatial consistency values from m1 versus from m2. Examples are shown in Figure 3H-J (metric values indicated in black-edged scatters in Figure 3E,F). The inferred cell ID-shuffled test data trajectories tend to have small steps and locate far away from both manifolds, having diminutive spatial consistency values. Compared with original test data trajectory, circular-shuffled data have smooth trajectories around the same area, with slightly smaller spatial consistency values and comparable step distances. Latent variables of time-shuffled test data has similar location as the original test data does, but are mostly taking big jumps across time. Just like the original test data, both circular-shuffled and time-shuffled test data have similarly high m1 contribution to spatial consistency and diminutive m2 contribution, suggesting the test data is repeating neural state encoding experience in maze1 rather than maze2. Recall that circular-shuffled data have scrambled coactivation patterns, which indicates that local neuronal firing rates can already distinguish animal contexts.

### 4.3 PBE decoding

In the previous two sections, it has been demonstrated that the model learned from training data during running can capture the neural state repetition in test data during running well. Next, we evaluated whether this running-state model could be used to estimate the internal state repetition in neural data during PBEs, which occurred when the animal had paused running in the exploration session.

First, the best time bin size for PBE data was searched. Both in the maze1 and maze2 exploration sessions, the 80^th^ percentile of 4 seconds’ travel distance is 86-87cm. Therefore, only place cell pairs with their place field distances less than 87cm are included in the subsequent analysis. It turns out that, for both maze sessions, the sum of significant (*p <* 0.01) positive Pearson correlation coefficient *r* between CCH during running and CCH during PBEs across cell pairs reached its peak when the PBE time bin size is 17 ms. Therefore, PBEs were binned into 17 ms time bins.

Next, in the latent space learned using running data from the two maze exploration sessions described in Section 4.2, we examined whether the inferred latent trajectories of binned PBE data could be identified as neural state repetitions. Firing rates of the tuning curve vectors were scaled by the time bin size ratio 17/500. Among 562 PBEs detected within the maze1 exploration session, we randomly selected 100 PBEs as test data. To evaluate the resulting latent trajectories, for each PBE, 200 surrogate data for each of the three shuffling types were generated and metric values were assessed.

The first question to ask about the PBE latent trajectories is the validity of neural state repetition. We consider a PBE to be a significant replay event, when its LLH value is larger than all of those of its circular-shuffled versions and its modified z-scored LLH is greater than 3. Moreover, a significant replay event should have a latent trajectory located closely to the tuning curve latent variable 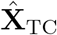 manifolds to guarantee the strength of the tuning curve constraint. Therefore, the average distance of the latent trajectory to its nearest neighbors identified during measuring the spatial consistency value is estimated. PBEs with average nearest neighbor distance larger than the standard deviation of 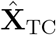 are excluded. Given that neuron firings of a replay are not exactly the time-compressed version of those during corresponding behavior, this criterion is a strict indicator of the ensemble activity pattern reactivation, and therefore, a strict indicator of the validity of being a replay event. Following the identification as a replay, z-scored LLH among its time-shuffled versions suggests the likeliness of being a *sequential* replay. Figure 4A shows the LLH values z-scored by circular-shuffled versions versus z-scored by time-shuffled versions, of the 100 selected PBEs and of 8 continuous running segments in maze1 for comparison. Each PBE is depicted as a dot, whose size stands for its spatial consistency value. 50 out of the selected 100 PBEs meet the replay criterion and are considered significant replay events, colored according to their z-scored LLH among time-shuffled versions. Insignificant PBEs are colored in gray.

**Figure 4:**
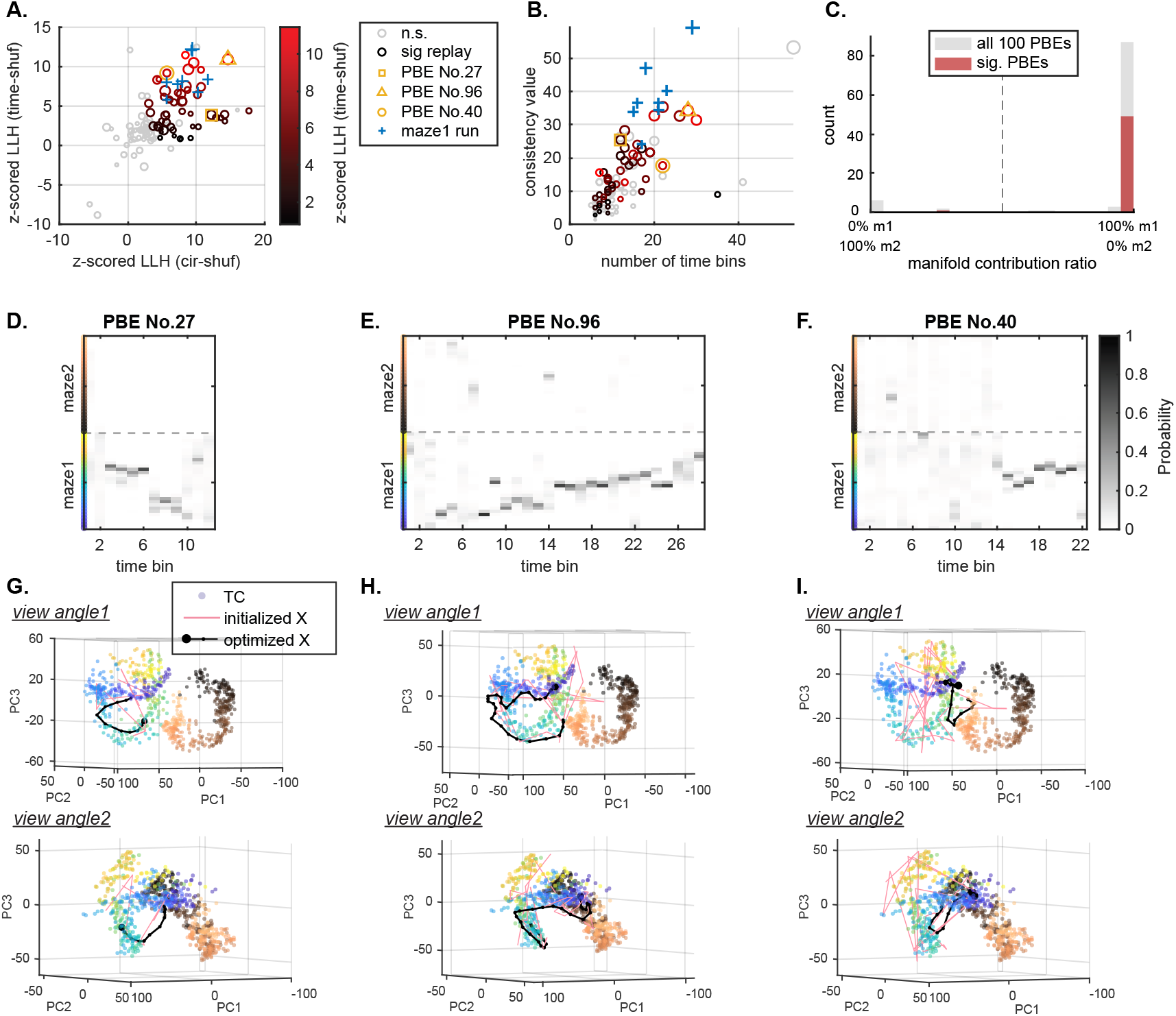
Inferring latent trajectories of PBE data in the P-GPLVM latent space learned from running data. **(A)** Modified z-scored LLH values among circular-shuffled versions versus among time-shuffled versions, of the 100 selected PBEs and of 8 continuous running segments in maze1 for comparison. The size of each circle indicates the spatial consistency value of the PBE. PBEs not meeting the replay criterion are in gray and PBEs identified as significant replay events are colored according to their z-scored LLH among time-shuffled versions. **(B)** Number of time bins of each test data versus their spatial consistency to the latent variable manifolds of tuning curve vector,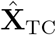. Dots are presented in the same way as in (A). **(C)** Histogram of ratio of manifold contribution to spatial consistency. Most of the PBEs are associated with maze1. **(D-F)** For comparison, we use traditional Bayesian decoding to render the posterior position distribution of the example PBEs (No.27, No.96, No.40) indicated in markers in (A-B). Corresponding context and position colors are indicated on the left side, same color code as Figure 3A. **(G-I)** Inferred latent trajectories of the example PBEs in (D-F) in the P-GPLVM latent space. While the initialized trajectories are jumpy, after iterations, the trajectories of the PBEs all converge to the maze1 manifold (same representation as in Figure 3G).

Next, the spatial distribution and dynamics features of each PBE were investigated. Since it had been shown that position information is encoded along the manifolds in latent space, for comparison, we estimated spatial tuning curves in the two running directions separately and used a Bayesian model to decode the PBE positions following the traditional method. Among those probable replay events, three example PBEs of different trajectory behaviors are highlighted by different markers in Figure 4A-B, whose Bayesian decoding results and inferred latent trajectories are visualized subsequently in Figure 4D-I. Figure 4B shows the number of time bins of each PBE and its spatial consistency to 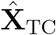. Visually, there is a trend that more time bins lead to larger spatial consistency values, which is intuitive because longer trajectory can travel through more area. PBEs in this trend indicate one-directional replays of experiences. For example, PBE No.96 in Figure 4E and H is a continuous replay of experience traveling from location in purple all the way to location in green, consistent with the Bayesian decoded results. PBEs with fewer time bins relative to those with comparable spatial consistency values (top left side of the chart) indicate that their trajectories might be of high speed or even have leaps. PBE No.27 in Figure 4D and G is shown as an example of this kind, whose Bayesian decoded results suggest a travel from location green to location purple with a leap in the middle. On the contrary, PBEs with more time bins relative to those with comparable spatial consistency values (bottom right side of the chart) indicate that their trajectories might be stationary or slightly oscillating. One example is PBE No.40 in Figure 4F and I, whose Bayesian decoded positions are first unfathomable (which might not be able to be decoded by a spatial decoder) and then travel around location green. Its latent trajectory at the first major part jiggled around location purple and then traveled to location green. Figure 4C shows the manifold contribution to spatial consistency value from m1 manifold versus from m2 manifold of each PBE. As expected, since the animal hadn’t experienced maze2 yet, almost all of the probable replay events belong to m1 manifold rather than to m2 manifold, indicating that they are replay events of maze1 experience.

Combining the observation of metric values, we can estimate the validity of neural state repetition, evaluate the pattern of this repetition, and infer the external variables of the replayed experience. *Note that we have not used position data in any of our analyses. This approach is the first presentation of a method for continuously decoding and identifying replay events without relying on external variables*.

Finally, with no need of assuming external variables that drive the neural activity or predetermining replay behavior patterns, the summary of replay behavior through out all sessions in this experiment can be revealed by applying the extended P-GPLVM model to all PBEs detected. Latent neural trajectories of all PBEs are inferred. For each PBE, 100 circular-shuffled versions were generated to for identifying significant replay events. PBE occurrence time vs. the ratio of manifold contribution to PBE spatial consistency value is shown in Figure 5 from pre session to post2 session in time order, where each PBE is depicted as a circle, whose size indicates its spatial consistency value to 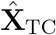. Circles on the top / bottom line means 100% of identified nearest neighbors for the latent neural trajectories are from m1 / m2 manifold, indicating affiliation with maze1 / maze2 context experience. Circles located in between suggest mixtures. PBEs identified as significant replay events are colored according to its z-scored LLH value among their circular-shuffled versions. In the pre rest session, identified significant replay events are comparably sparse and have small spatial consistency values. In maze1 and maze2 exploration sessions, as expected, almost all the significant replay events are associated with their current contexts and have comparably larger consistency and z-scored LLH values, which indicates long and sequential experience replay. In post1 rest session, the majority of significant replay events are associated with maze1 context and most of them have small consistency values. Those replay events became less associated with maze1 as the rest time increased. In post2 rest session, replay events seem to be more frequent and most of them are associated with maze2 context.

**Figure 5:**
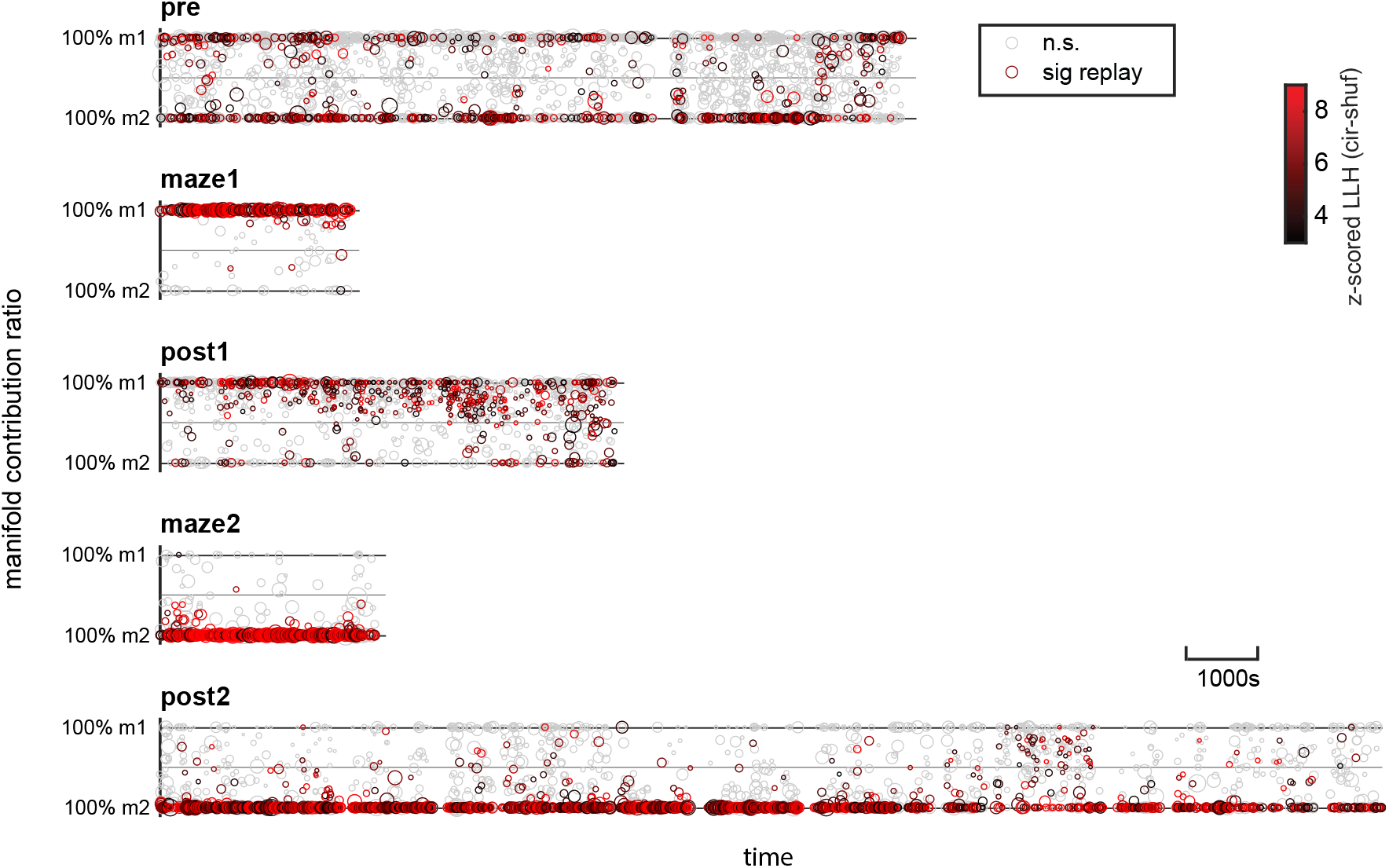
PBE occurrence time vs. ratio of manifold contribution for PBE latent trajectories in all sessions. The top solid line indicates 100% of the trajectory nearest neighbors are from maze1 manifold (m1). The bottom solid line indicated 100% of the nearest neighbors are from maze2 manifold (m2). Each circle indicate one detected PBE. Significant replay events are colored in black to light red according to their z-scored LLH values among their circular-shuffled versions.

## Conclusion

As the results have shown, without referring to any external variables, P-GPLVM is a powerful tool to capture nonlinear neural population dynamics in the hippocampus by discovering the low-dimensional structure underlying the neural activity, which reveals important encoded external variables such as the number of contexts and the topology of animal spatial behavior (i.e. position). In this paper, this model is extended to leverage those advantages in the neural decoding scenario by enabling the constrained latent variable inference of new neural data and proposing a family of analyses for result evaluations. This constrained inference approach requires much less computational cost than relearning the model with training data combining old and new data. This extended model is flexible that, for new neural data either during running or during PBEs, neural trajectories can be inferred in the latent space learned from training data during running and internal neural state repetition can be evaluated. External variables can then be decoded based on the external experiences corresponding to the repeated neural states. Metrics are defined that for the first time enable the identification of continuously-decoded replay using a model trained only with neural activity (and refine this definition to allow for both sequential and non-sequential events).

## Acknowledgments

This research was supported by R01NS115233.

